# Human Galectin-9 Promotes the Expansion of HIV Reservoirs *in vivo* in Humanized Mice

**DOI:** 10.1101/2022.07.19.500638

**Authors:** Zhe Yuan, Leila B. Giron, Colin Hart, Akwasi Gyampoh, Jane Koshy, Kai Ying Hong, Toshiro Niki, Thomas A. Premeaux, Lishomwa C. Ndhlovu, Luis J Montaner, Mohamed Abdel-Mohsen

## Abstract

**Objective:** The human endogenous β-galactoside-binding protein Galectin-9 (Gal-9) reactivates latently HIV-infected cells, which may allow for immune-mediated clearance of these cells. However, Gal-9 also activates T cell Receptor (TCR) signaling pathways, which could negatively affect HIV persistence by promoting T cell expansion and chronic activation/exhaustion. This potential “double-edged sword” effect of Gal-9 during HIV infection raises the question of the overall beneficial versus detrimental impact of Gal-9 on HIV persistence *in vivo*.

**Design:** We used the BLT (bone marrow, liver, thymus) humanized mouse model to evaluate the overall impact of Gal-9 on HIV persistence *in vivo* during antiretroviral therapy (ART).

**Methods:** Two independent cohorts of BLT mice with high human immune reconstitution were infected with HIV, placed on ART, and then treated with either recombinant human Gal-9 or PBS during ART suppression. Plasma viral loads and levels of tissue-associated HIV DNA and RNA were measured by qPCR. Markers of T cell activation/exhaustion were measured by flow cytometry, and plasma markers of inflammation were measured by multiplex cytokine arrays.

**Results:** Gal-9 treatment was tolerable in ART-suppressed humanized mice and did not significantly induce plasma markers of inflammation or T cell markers of activation/exhaustion. However, Gal-9 treatment during ART significantly increased levels of tissue-associated HIV DNA and RNA compared to controls (*P*=0.0007 and *P*=0.011, respectively, for cohort I and *P*=0.002 and *P*=0.005, respectively, for cohort II).

**Conclusions:** Our study highlights the overall adverse effects of Gal-9 on HIV persistence and the potential need to block Gal-9 interactions during ART-suppressed HIV infection.

## INTRODUCTION

The persistence of HIV latently-infected cells in blood and tissues despite effective antiretroviral drug therapy (ART) remains a barrier to a cure for HIV infection ^[1]^. Persistent latently-infected cells do not express enough viral antigens to be eliminated by the immune system, and different strategies to reactivate these cells have been proposed, including the use of the human lectin Galectin-9 (Gal-9) ^[2, 3]^. However, Gal-9 modulates HIV transcription through activating the T-cell receptor (TCR)-downstream ERK and CREB signaling pathways ^[4]^ and this Gal-9-mediated activation can lead to several undesirable effects due to T cell expansion, activation, and/or exhaustion. Indeed, recent studies showed that Gal-9, which is rapidly and sustainably elevated during HIV infection ^[5]^, may contribute to the state of chronic immune activation and inflammation during HIV infection ^[6-8]^. This potential “double-edged sword” of Gal-9 during HIV infection raises the question on how the potentiation of reactivating latent infection and the undesirable outcomes on T cell effector function due to Gal-9 effects HIV persistence *in vivo*. Thus, evaluating the overall impact of Gal-9 on HIV persistence requires a relevant *in vivo* animal model of HIV infection during ART suppression. In this short report, we investigate the impact of Gal-9 in a humanized mouse model of HIV infection (the bone marrow-liver-thymus humanized (BLT) mouse model of HIV latency) to determine the overall beneficial versus detrimental impact of Gal-9 on HIV persistence *in vivo*.

## METHODS

### Generation of the BLT mice

BLT mice were generated as previously described ^[9, 10]^, in accordance with The Wistar Institute Animal Care and Research Committee regulations. Briefly, 6-8 weeks old female NSG (NOD.Cg-Prkdcscid Il2rgtm1Wjl/SzJ, Jackson Laboratory) mice were pretreated with busulfan at 30mg/kg and were then implanted with human fetal thymic tissue fragments and fetal liver tissue fragments under the murine renal capsule. Following the surgery, mice were injected via the tail vein with CD34^+^ hematopoietic stem cells isolated from human fetal liver tissues. Human fetal liver and thymus tissues were procured from Advanced Bioscience Resources (Alameda, CA). Twelve weeks post-surgery, human immune cell reconstitution in peripheral blood was determined by Symphony flow cytometer (BD Biosciences, San Jose, CA) using the following antibodies: mCD45-AF700, hCD45-FITC, hCD3-BUV805, hCD4-BUV395, hCD8-PerCP-Cy5.5 and Fixable Viability Stain 510 (BD Biosciences, San Jose, CA). Data were analyzed with FlowJo (FlowJo LLC, Ashland, OR).

### HIV infection, ART suppression, and Gal-9 treatment

BLT mice were randomly divided into two groups and were infected intravenously (IV) with 1×10^4^ TCID_50_ of HIV_SUMA_. Peripheral blood was collected weekly for plasma viral load assay. Two weeks post-infection, mice were placed on a diet combined with ART (1,500mg/kg Emtricitabine, 1,560mg/kg Tenofovir-Disoproxil-Fumarate, and 600mg/kg Raltegravir). Five weeks post-ART, mice were treated with phosphate-buffered saline (PBS) control or 2 mg/kg recombinant Gal-9 for two weeks (intraperitoneal (IP) injections every other day; seven doses) during ART suppression. Mice were then euthanized, and blood and tissues were collected.

### Measuring plasma viral load by qPCR

Plasma viral loads were measured as previously described ^[9, 10]^. Briefly, viral RNA was extracted using the QIAamp Viral RNA Mini kit (Qiagen). Plasma viral load was measured using qRT-PCR on a C1000 Thermal Cycler and the CFX96 Real-Time system (Bio-Rad) with 100 copies/ml limit of detection.

### Measuring cell-associated HIV DNA and RNA by qPCR

Single cells are isolated with gentleMACS™ Octo Dissociator (San Diego, CA). DNA and RNA were extracted using AllPrep DNA/RNA/miRNA Universal Kit (Qiagen, catalog # 80224). Cell-associated HIV DNA was measured by qPCR using LTR-specific primers F522-43 (5’ GCC TCA ATA AAG CTT GCC TTG A 3’; HXB2522–543) and R626-43 (5’ GGG CGC CAC TGC TAG AGA 3’; 626–643) coupled with a FAM-BQ probe (5’ CCA GAG TCA CAC AAC AGA CGG GCA CA 3). Cell count was estimated by qPCR using human copy number reference assay TERT (Applied Biosystems, cat# 4403316). Cell-associated HIV RNA was measured using the same primers in a one-step qPCR using TaqMan® RNA to Ct™ 1 Step kit (Applied Biosystems, catalog # 4392656). Cell counts were determined by qPCR using human RPLP0 (Applied Biosystems, catalog # 4310879E).

### Measuring markers of T cell activation and exhaustion by flow cytometry

Cell suspension was stained with the following antibodies: CD45-AL700, CD8-FITC, CD38-APC, CD25-PerCP-Cy5.5, HLA-DR-APC-H7, CD4-V450, PD-1-PE, CD3-PE-CF594, and CD69-PE-Cy7. Data were collected on a BD Biosciences LSRII flow cytometer (gating strategy is in **Supplementary Figure 1**).

### Measuring plasma markers of inflammation

Markers of inflammation were measured using U-PLEX Biomarker Group 1 Assay from Meso Scale Discovery (MSD catalog # K15067L-2).

### Statistical analysis

Mann–Whitney tests were used for the two-group comparisons in Figures 1B, G, H, and unpaired t-tests were used for the two-group comparisons in Figure 2. Statistical analyses were performed using Prism 9.0 (GraphPad).

**Figure 1.**
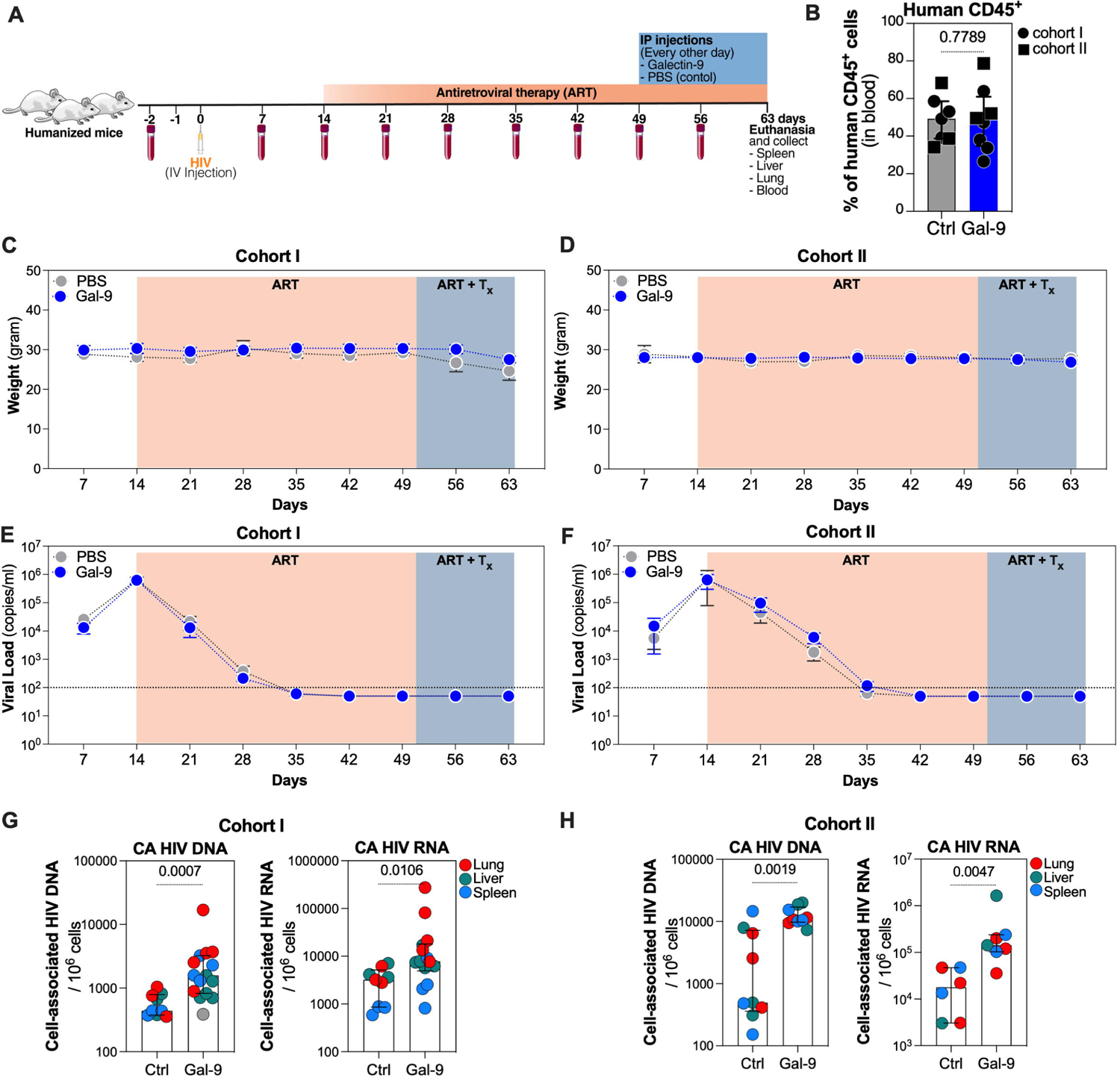
Gal-9 treatment is tolerable *in vivo* but increases levels of tissue-associated HIV DNA and RNA during ART-suppressed HIV infection. **(A)** a schematic overview of the study design. **(B)** percentage of human CD45^+^ cells measured in the peripheral blood of BLT mice. No difference in levels of human immune reconstitution between mice treated with Gal-9 versus mice treated with PBS control. Mann Whitney t test. Median and interquartile range (IQR) are displayed. (**C-D**) mice weight over time in cohort I (C) and cohort II (D). **(E-F)** plasma viral load measured by ddPCR over time in cohort I (E) and cohort II (F). **(G-H)** Gal-9 treatment-induced levels of tissue-associated HIV DNA and HIV RNA (measured by qPCR) than controls in both cohort I (G) and cohort II (H). Data shown are only from tissues with detectable levels of cell-associated HIV DNA or RNA. Mann Whitney t-tests. Median and interquartile range (IQR) are displayed.

**Figure 2.**
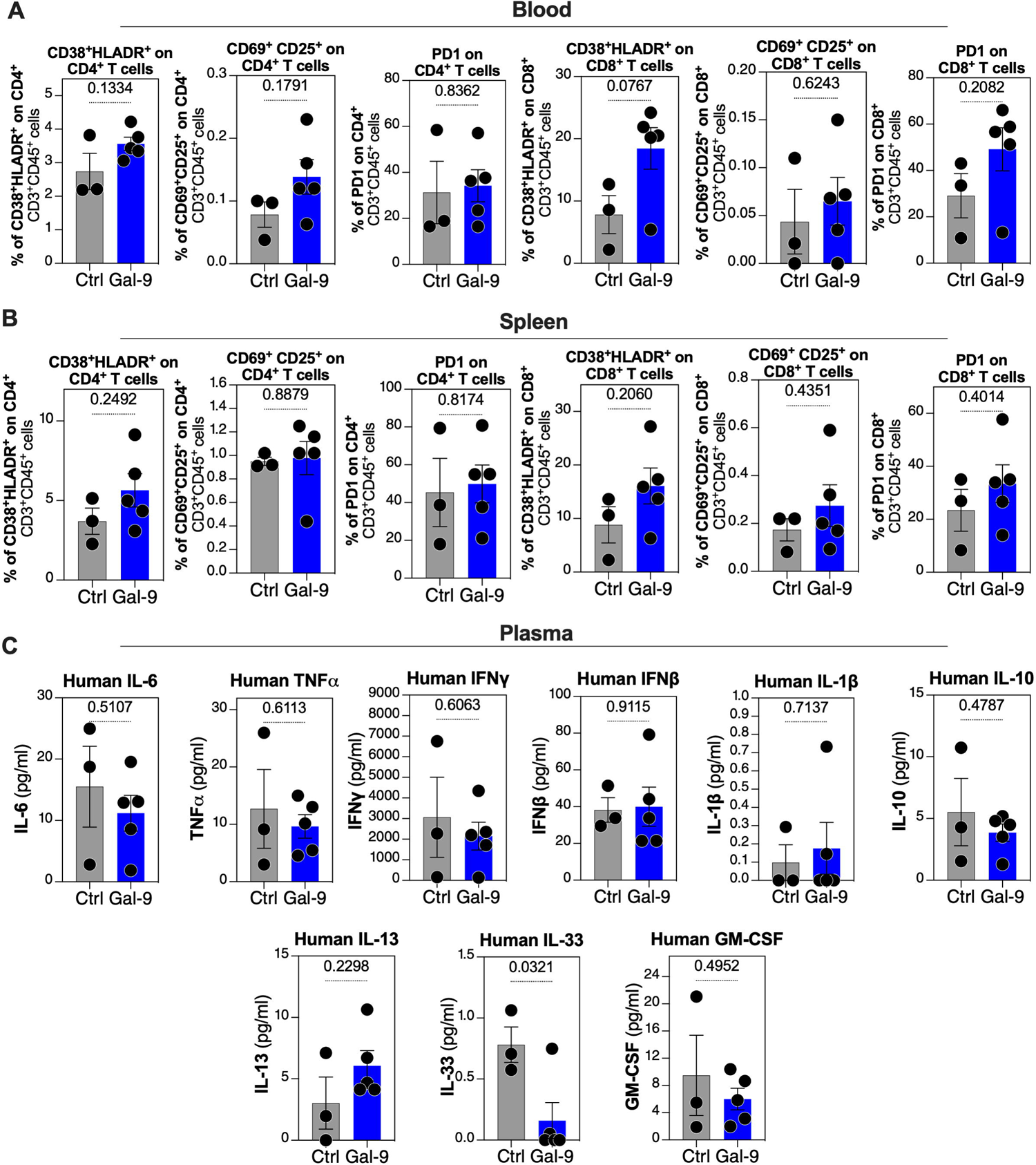
Gal-9 treatment does not significantly induce markers of inflammation of T cell activation. **(A-B)** impact of Gal-9 treatment on markers of CD4^+^ and CD8^+^ T cell activation (co-expression of CD38 and HLA-DR or the co-expression of CD69 and CD25 activation markers) and exhaustion (PD-1 expression) measured by flow cytometry in the blood (A) and spleen (B). Unpaired t-tests. The mean and standard mean of error (SEM) are displayed. **(C)** Impact of Gal-9 treatment on plasma levels of markers of systemic inflammation measured by multiplex arrays. The mean and standard mean of error (SEM) are displayed.

## RESULTS

### Gal-9 treatment is tolerable *in vivo* but increases levels of tissue-associated HIV DNA and RNA during ART

We generated two independent cohorts of humanized mice (n=9 for cohort I and n=6 for cohort II) (**Fig 1A**) and achieved high levels of immune reconstitution as measured by the percentage of human CD45^+^ cells in blood (**Fig. 1B**). BLT mice were infected with HIV (HIV_SUMA_ transmitted/founder virus) and then placed on ART two weeks post-infection. During ART suppression, mice were treated with either PBS or 2 mg/kg recombinant human Gal-9 for two weeks (every other day; seven doses). Mice were then euthanized, and blood and tissues (liver, spleen, and lung) were collected (**Fig. 1A**). Gal-9 treatment did not affect the weight of the mice from both cohorts compared to controls **(Fig. 2C-D)**, suggesting this concentration was generally tolerable in ART-suppressed HIV-infected BLT mice. Plasma HIV viral loads shown in **Fig. 1E-F** indicate that, in both cohorts, the infection was successful and resulted in high viral loads by the second week post-infection and that ART suppressed the virus to below the limit of detection. We have not observed viral blips in the Gal-9 treated mice to suggest any significant viral reversal effects. We next examined cell-associated HIV RNA and DNA levels in the liver, lung, and spleen. We found that Gal-9 treatment increased the levels of tissue-associated HIV DNA (*P*=0.0007) and RNA (*P*=0.0106) (**Fig. 1G**) in cohort I. Consistently, in cohort II, Gal-9 increased tissue-associated HIV DNA (*P*=0.0019) and RNA (*P*=0.0047) (**Fig. 1H**). Collectively, these data suggest that the overall impact of Gal-9 on HIV persistence is negative, and that Gal-9 expands the levels of tissue-associated HIV reservoirs during ART.

### Gal-9 treatment does not significantly induce markers of inflammation of T cell activation

Next, we examined whether Gal-9 exhibits its adverse effects on HIV persistence by inducing T cell activation/exhaustion or systemic inflammation. We first examined markers of CD4^+^ and CD8^+^ T cell activation/exhaustion in blood and spleen (collected at the time of mice euthanization from cohort I). Gal-9 treatment did not significantly induce T cell activation (as measured by the co-expression of CD38 and HLA-DR or the co-expression of CD69 and CD25 activation markers) or exhaustion (as measured by PD-1 expression) in blood (**Fig. 2A**) or tissues (**Fig. 2B**). However, a trend toward an increase in CD8^+^ T cell expression of CD38 and HLA-DR was observed in the blood in the Gal-9 treated mice compared to controls (p=0.0767, **Fig. 2A**). Next, we examined the plasma levels of several cytokines. While Gal-9 treatment reduced the levels of IL-33 (p=0.0321), it did not significantly alter the systemic levels of several cytokines involved in inflammation (**Fig. 2C**). These data suggest that Gal-9 does not induce levels of cell-associated HIV DNA and RNA by inducing generalized T cell activation or systemic inflammation.

## DISCUSSION

A comprehensive understanding of the overall impact of viral and host factors that modulate HIV persistence is critical to develop curative strategies for HIV. Gal-9 is one of the endogenous host immune-modulatory factors that has been recently associated with opposing effects on HIV infection ^[5, 11, 12]^. Several studies have highlighted the potential beneficial effects of Gal-9 during HIV infection. First, Gal-9 renders CD4^+^ T cells less susceptible to HIV infection via induction of the host restriction factor cyclin-dependent kinase inhibitor 1 (p21) ^[13]^. Second, recombinant Gal-9 induces HIV transcription and reverses HIV latency *in vitro* and *ex vivo*. This ability of Gal-9 to induce latent HIV transcription suggested that it could be considered within the “shock and kill” HIV eradication framework ^[2, 3]^. However, on the other hand, several other studies have highlighted the detrimental effects of Gal-9 during HIV infection. First, endogenous Gal-9 is rapidly increased after HIV infection ^[5]^ and do not return to normal after suppressive ART ^[5]^. These persistent elevated Gal-9 levels are associated with higher HIV transcription *in vivo* in plasma of HIV-infected ART-suppressed individuals ^[2]^, and several studies suggest may contribute to the state of chronic immune activation and inflammation during HIV infection ^[6-8]^. Second, Gal-9 increases HIV entry by inducing the CD4^+^ T cell-surface concentration of protein disulfide isomerase (PDI) ^[14]^. Lastly, Gal-9 modulates HIV transcription by activating the TCR-downstream signaling pathways *in vitro* ^[4]^. Due to these pleiotropic effects of Gal-9 on HIV, we sought to evaluate whether the overall impact of Gal-9 on HIV persistence is beneficial or detrimental *in vivo*.

Using the BLT humanized mouse model of HIV infection, we found that Gal-9 can directly expand HIV reservoirs and exhibit overall adverse effects on HIV persistence. These effects would limit the potential use of Gal-9 to reduce viral reservoirs by itself. However, future studies will be needed to investigate whether Gal-9 together with added immune effector strategies would positively affect HIV persistence. Future studies will also be needed using larger animal models with a more intact immune system compared to the BLT humanized mouse model. Our data also show that the Gal-9-mediated expansion of HIV reservoirs is not linked to elevated T cell activation or systemic inflammation. However, Gal-9 can induce T cell proliferation ^[2]^, which might be linked to its modulation of HIV persistence. Furthermore, Gal-9 can impact many aspects of immune responses relevant to persistence. For example, Gal-9 exhibits several immunosuppressive activities ^[15-23]^, including the ability to increase the function of regulatory T cells (T-regs)^[23]^ and to impair natural killer (NK) cells’ cytotoxicity ^[24]^. The potential link between Gal-9-mediated impact on T cell proliferation or immune functions and its ability to impact HIV reservoir size warrants a broader investigation.

Endogenous Gal-9 is highly abundant *in vivo*, especially during HIV infection^[5]^, and its sustained levels have been associated, during ART-suppressed HIV infection, with the state of chronic inflammation and immune activation that is central to the development of several HIV-associated co-morbidities ^[25-32]^. Therefore, clarifying the mechanistic underpinnings of the overall adverse effects of elevated Gal-9 during ART-suppressed HIV infection may lead to the development of interventions to target Gal-9 (such as anti-Gal-9 antibodies and small molecule inhibitors targeting Gal-9 ^[33-35]^) to improve immune functionality, reduce inflammation-associated co-morbidities, and reduce levels of HIV persistence, in the setting of viral suppression by ART.

## SUPPLEMENTARY MATERIALS

**Supplementary Figure 1. A gating strategy [Leila]**. A representative example of the gating strategy used for the phenotyping of markers of T cell activation and exhaustion in Figure 2.

## AUTHOR CONTRIBUTIONS

Z.Y, L.B.G, T.N, L.C.N, L.J.M, and M.A-M designed the experiments. Z.Y, L.B.G, C.H, A.G, J.K, and K.Y.H carried out the experiments. Z.Y, L.B.G, L.J.M, M.A-M analyzed and interpreted data. Z.Y, L.B.G, L.J.M, M.A-M wrote the manuscript, and all authors edited it.

