## Supplementary figures and images for "Human Galectin-9 Promotes the Expansion of HIV Reservoirs *in vivo* in Humanized Mice"

### Supplementary Figure 1

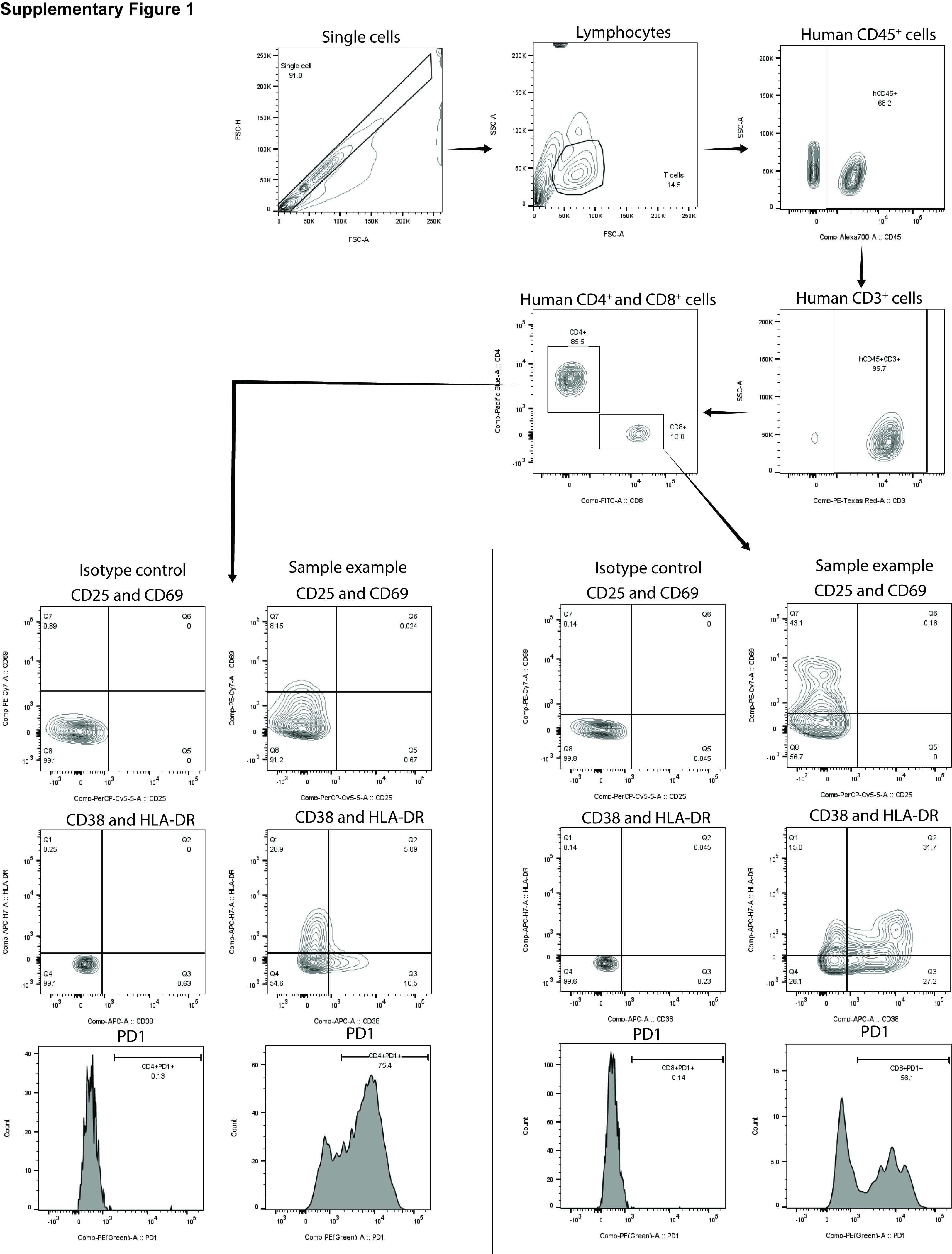
